# Dietary ethanol affects feeding behaviour, flight performance, and mating success in a fruit-feeding butterfly

**DOI:** 10.64898/2026.01.13.699346

**Authors:** Nuno Dias Osório, Emilia Potselueva, Sridhar Halali, Patrícia Beldade, Erik van Bergen

**Affiliations:** Department of Biology, Faculty of Sciences, University of Lisbon (FCUL), Lisbon, Portugal; Biodiversity and Evolution Unit, Department of Biology, Lund University, Lund, Sweden; Centre for Ecology, Evolution and Environmental Changes (CE3C) & Global Change and Sustainability Institute (CHANGE), Faculty of Sciences, University of Lisbon (FCUL), Lisbon, Portugal

**Author notes:** **Author contributions.** N.D.O., E.P., S.H. and E.vB. conceived and designed the study. N.D.O. and E.P. conducted the experiments and collected the data. E.vB. performed the data analysis and prepared the figures. P.B. provided all laboratory and research equipment. E.vB. wrote the original draft of the manuscript, with support from P.B. Portions of the text were revised with assistance from OpenAI’s ChatGPT (GPT-4, accessed between August and December 2025) to improve clarity and style. All authors critically reviewed and edited the manuscript and approved the final version. **Data availability statement.** Data will be made available upon acceptance of the manuscript (e.g. Dryad Digital Repository).

**Keywords:** frugivory, fermentation, alcohol, insect behaviour, tropical ecology, *Bicyclus anynana*

## Abstract

1. The evolution of fleshy fruits around 124 million years ago introduced a nutritious resource and drove shifts toward fruit-feeding across the animal kingdom, leading to a variety of adaptations for locating and exploiting this resource. Fruits are inherently dynamic, changing in physical, chemical, and biological properties as they progress from development through ripening to decay.
2. Fruit fermentation produces ethanol, a volatile compound that is simultaneously caloric, intoxicating, and potentially noxious. While the immediate and long-term effects of dietary ethanol are well documented in mammals, and to some extent in insects such as *Drosophila*, very little is known about its impact on frugivorous butterflies.
3. Although ethanol can serve as a foraging cue for frugivorous butterflies, its ingestion may impair performance and reduce fitness. Here, we examine how dietary ethanol affects ecologically and reproductively relevant behaviours in the Afrotropical butterfly *Bicyclus anynana*, a species that feeds on fermenting fruits in the wild.
4. We found that butterflies exposed to ethanol-supplemented diets initiated feeding more rapidly, but also spent less time feeding, and they required more stimuli to sustain flight. Moreover, feeding behaviour differed between males and females and between mated and virgin individuals, while flight performance differed between sexes but was not affected by mating status.
5. Given the involvement of flight in male courtship, we also tested the hypothesis that negative effects of ethanol consumption on flight would impair courtship. We found that males fed on ethanol-supplemented diets—particularly young ones—had reduced mating success and took longer to secure a mating.
6. Together, our findings demonstrate that ethanol consumption affects behaviours closely tied to survival and fitness in *B. anynana*, suggesting that exposure to fermenting resources may impose selective pressures on physiology, behaviour, and reproductive strategies in frugivorous insects.

## Introduction

The strategies organisms use to obtain and process nutritional resources are widely recognized as major drivers of evolutionary change (Nosil, 2012; Schluter, 2000). As species expand or shift the range of resources they exploit, natural selection favours morphological, physiological, and behavioural traits that enhance their ability to acquire, process, and utilise these resources. This is particularly evident in plant-feeding animals, which exploit diverse tissues (e.g., leaves, stems, seeds, fruits, nectar, and roots), each presenting distinct nutritional, physical, and chemical challenges. These challenges impose strong selective pressures, often resulting in specialized feeding morphologies (Grant & Grant, 2008; Seehausen, 2006), digestive adaptations (Hofmann, 1989; Mackie, 2002), and detoxification mechanisms (Edger et al., 2015; Ratzka et al., 2002), which drive adaptive divergence among lineages. Classic examples of such resource-driven diversification include the radiation of Darwin’s finches on the Galápagos Islands, where variation in food types resulted in an extraordinary array of beak shapes and feeding strategies (Grant & Grant, 2008; Lamichhaney et al., 2015), and the explosive diversification of East African cichlids, whose jaw morphologies reflect finely tuned adaptations to different dietary niches (Brawand et al., 2014; Seehausen, 2006).

Since the evolution of fleshy, sugar-rich fruits by angiosperms, approximately 124 million years ago (Eriksson, 2016; Tiffney, 1984), a wide range of animals have evolved specialized behavioural, sensory, physiological, and morphological adaptations that enable them to efficiently locate and consume these nutrient-rich resources (Herrera, 2002; Jordano, 2000). These adaptations are tuned to the biological, chemical, and physical properties of fruit, which change during development, ripening, and subsequent rotting. Frugivorous birds, bats and primates typically rely on visual cues such as colour change to detect ripe and overripe fruits (Nevo et al., 2018; Schaefer et al., 2004), while other frugivores use volatile compounds produced during fermentation to locate overripe or decaying fruit (Becher et al., 2012; Utrio & Eriksson, 1977). During ripening and decay, naturally occurring microorganisms convert sugars into ethanol through anaerobic fermentation, a process that supports their energy needs and/or suppresses the growth of ethanol-intolerant competitors (Dashko et al., 2014; Piskur et al., 2006). Fermentation also breaks down plant cell walls and complex carbohydrates into monosaccharides, enabling frugivores to extract nutrients more efficiently from ripe or overripe fruits.

Animals can use ethanol and other byproducts from the fermentation process as olfactory cues to optimize foraging efficiency (Dudley, 2004). For example, African elephants assess the ethanol concentration of marula fruits to preferentially select those with the highest sugar levels (Nevo et al., 2020). Similarly, fruit flies (*Drosophila* spp.) are well-documented to detect ethanol and other fermentation volatiles, which guide them to suitable substrates for feeding and oviposition on overripe or fermenting fruits (Becher et al., 2012; Devineni & Heberlein, 2009; Hodgkison et al., 2013). While ethanol can act as a foraging cue, its consumption can have detrimental effects, especially in organisms lacking effective detoxification pathways. Exposure to ethanol depresses central nervous system activity in animals, and may impair motor coordination, reaction time, and decision-making—even at low doses (Bowland et al., 2024; Devineni & Heberlein, 2013). For instance, ethanol consumption has been shown to impair flight performance and echolocation in captive Egyptian fruit bats (Sánchez et al., 2010; also see Bowland et al., 2024), as well as locomotor behaviour (Maze et al., 2006) and foraging decision-making (Abramson et al., 2005) in honeybees (*Apis mellifera*). In the latter case, ethanol-exposed individuals choose flowers at random rather than selectively targeting those offering higher energetic rewards, as is typical for the species (Abramson et al., 2005).

To cope with the adverse effects of dietary ethanol, many frugivorous animals—or those exposed to ethanol through sources like fermenting nectar—have evolved specialized adaptations that enable efficient alcohol metabolism (Ashburner, 1998; Chambers, 1988). These adaptations include alcohol dehydrogenase (ADH) and aldehyde dehydrogenase (ALDH), enzymes that catalyse the metabolic breakdown of ethanol into acetate, a less harmful compound used in cellular energy pathways (Fry, 2014; Geer et al., 1993). Notably, studies in fruit-feeding insects (e.g., *Drosophila*) and hornets have shown that duplications of the *Alcohol dehydrogenase* (*Adh*) gene are associated with adaptation to alcoholic niches (Atrian et al., 1998; Bouchebti et al., 2024). Similarly, dietary specialization on fruit or nectar in mammals intensified selective pressure on genes involved in ethanol metabolism, as well as on their catalytic activity (Carrigan et al., 2015; Janiak et al., 2020). Despite the presence of effective detoxification mechanisms, consumption of ethanol-containing food can still impair short-term performance and can reduce overall fitness by compromising survival, reproductive success, or competitive ability (Bowland et al., 2024).

Both immediate and long-term effects of ethanol are well-documented in mammals (Adamson et al., 2017), including humans (Le Daré et al., 2019). In contrast, apart from *Drosophila* species (Keesey et al., 2025; Wang & Althoff, 2019), such evidence remains scarce for frugivorous or nectivorous species in which ethanol intake may simply be a by-product of their niche specialization (but see Bowland et al., 2024). Indeed, these effects are relatively understudied in fruit-feeding butterflies, in spite of their ecological success. Here, we investigate the effects of ethanol on behaviours closely linked to survival and reproductive success in the tropical butterfly *B. anynana* (Brakefield et al., 2009). This species inhabits seasonal environments in East Africa (Aduse-Poku et al., 2022), where larvae feed on grasses (*Poaceae*) (van Bergen et al., 2016) and adults on fallen and decaying fruit, including that of *Ficus* trees (Larsen, 1991). Consequently, *B. anynana* is likely to be exposed to ethanol in its natural habitat, and fermented bait is commonly used to attract individuals in the wild (Mallick et al., 2024; Nokelainen et al., 2016). Moreover, as many as five putative *Adh* genes have been identified in this species (Conceição et al., 2011; Nowell et al., 2017).

While ethanol-supplemented diets had been shown to increase fecundity and survival in *B. anynana* females (Beaulieu et al., 2017), much less is known about how dietary ethanol affects key behavioural traits such as feeding behaviour, flight performance, and male mating success. The first part of our study focused on how diet-associated ethanol influences feeding and flight. Because ethanol is volatile, potentially intoxicating, and caloric, we hypothesized that ethanol consumption would reduce the time required for butterflies to locate food, the duration of feeding, and their flight performance. We found that diet-associated ethanol caused butterflies to initiate feeding faster, feed for shorter periods, and require more external stimuli to sustain flight. Moreover, given physiological and energetic differences associated with sex and mating status, we kept track of these variables. We found sex affected both feeding and flight behaviour, while mating status only affected feeding. The second part of our study focused on the effects of ethanol consumption on male mating success. Courtship in *B. anynana* involves males pursuing females through frequent take-offs and landings, as well as ritualized wing movements (Balmer et al., 2018; van Bergen et al., 2013). Our finding that ethanol impaired flight performance led us to hypothesize that it could also affect male mating success. We found that males fed on ethanol-supplemented versus control diets had reduced mating success (albeit only seen for younger males) and took longer to secure a mating. Taken together, our results reveal clear effects of ethanol on traits likely to matter to fitness. We discuss these results in the light of their potential ecological and evolutionary significance.

## Materials and Methods

### Animals and rearing conditions

*Bicyclus anynana* (Butler, 1879) is a butterfly species that inhabits seasonal environments across East Africa, with a geographic distribution ranging from Ethiopia to the Northern-most provinces of South Africa (Larsen, 1991; Windig, 1994). The butterflies used in this study originated from a captive population maintained at the University of Lisbon. This population was initially established from more than 80 gravid females collected at a single site in Nkhata Bay, Malawi (Brakefield et al., 2009), and has been maintained at large population sizes. All individuals were reared in a climate-controlled chamber set to 27 °C, 65% relative humidity, and a 12:12 light–dark cycle, mimicking environmental conditions these insects encounter during the wet season (van Bergen et al., 2016; Windig, 1994). Eggs were collected from the stock population every week to maintain a continuous supply of adult butterflies. Larvae were reared on young maize plants (*Zea mays*) in large cages at a density of 200 individuals with *ad libitum* feeding. Pupae were collected from the rearing cages and transferred to individual containers where adult emergence was monitored daily during weekdays. Upon emergence, adults were briefly anesthetized using CO₂ pads, sexed using morphological characters, and marked on both hindwings with a unique alpha-numeric identifier. Marked individuals were subsequently kept in same-sex hanging cages and fed using Petri dishes containing water-soaked cotton and banana slices.

### Mating and dietary ethanol treatments

After butterfly emergence and marking, individuals were randomly assigned to two mating treatments; half of the individuals of each sex were allowed to mate, while the other half remained unmated (see Figure S1 for a schematic representation). Mating pairs were isolated, and after mating, individuals were transferred to separate same-sex hanging cages (diameter: 30 cm; height: 40 cm). This resulted in four cages representing the following groups: unmated males, unmated females, mated males, and mated females. Following this, food was removed from all cages for approximately 72 hours, leaving only a Petri dish with water-soaked cotton to prevent butterfly dehydration. After fasting, butterflies from each cage were further divided into two groups of five individuals and randomly assigned one of two dietary treatments: a control diet consisting of mashed banana (90% of final volume) and water (10% of final volume), or an experimental diet containing mashed banana (90%) and azeotropic ethanol (10%; LabChem, Laborspirit Ltd). Since azeotropic ethanol is 96% ethanol by volume, the final ethanol concentration in the experimental diet was 9.6% v/v.

### Feeding and flight behaviour assays

At the beginning of each behavioural assay, a voice memo was initiated to document all relevant events. We recorded the time the food treatment was introduced into each cage and, using the unique alpha-numeric wing markings, the time each individual began feeding; together, these times defined the feeding latency (in seconds). We also recorded the interval between the observed start and end of feeding for each individual, which we defined as the total feeding time (in seconds). Feeding was identified by a butterfly sitting on the food and later leaving the food, rather than by continuous monitoring of proboscis extension.

Once individuals stopped feeding, they were each placed individually in an empty cage and subjected to a 1-minute forced flight test, following established protocols (Saastamoinen et al., 2010; van Bergen et al., 2013). Following brief acclimatisation, the netting cage was gently tapped to induce flight, and each time the butterfly alighted, it was immediately forced to take off again. Individuals were forced to take off at a maximum rate of one tap per second, corresponding to an upper limit of 60 taps per assay. The total number of taps was used to measure flight performance, with more taps indicating reduced flight endurance (see statistical analysis section). A total of 200 individuals were assayed across five replicate blocks.

### Male mating success assays

All individuals used in this assay were virgins at the time of the start of the assay. Males were randomly allocated to one of two cages, with carefully controlled age distributions to ensure comparability across experimental cohorts, while all females were maintained together in a large adult cage (see Figure S1). Prior to each mating assay, all males were deprived of food for 72 hours (see above; all females were fed *ad libitum* during this period). On the morning of the mating assay, both male cohorts (N≈10) were randomly assigned one of two dietary treatments, as described above: an experimental diet (banana mash with 10% azeotropic ethanol) or a control diet (banana mash with 10% water). After the males completed feeding, the Petri dishes containing the food were removed, and virgin females were added to a 1:1 sex ratio in each mating cage. From that point onwards, formation of mating pairs was observed continuously for 120 minutes. Once a pair was formed, the time of mating and the corresponding male wing marking were recorded before the pair was removed from the cage. Mating latency (in seconds) was defined as the time between the introduction of the females into the mating cage and the formation of the pair. At the end of the observation period, the unique identifiers of all unmated males were also documented. A total of 295 males, between three and ten days old, were assayed across 16 replicate blocks.

### Statistical analyses

All statistical analyses were conducted in R, v4.5.0 (R Core Team, 2025). Time variables (i.e. feeding latency, total feeding time, and mating latency) were initially analysed using generalized linear mixed-effects models (GLMMs) with the *glmmTMB* package in R (McGillycuddy et al., 2025), using a Gamma error distribution with a log link function, as is appropriate for modelling positively skewed continuous data (Ng & Cribbie, 2017). The attempt to model feeding latency using this strategy revealed significant overdispersion and violations of model assumptions. Consequently, this response variable was log-transformed and analysed using a GLMM with a Gaussian distribution.

Flight performance data (i.e. the number of times the butterfly was forced to take off over 60 seconds, with an upper limit of one ‘tap’ per second) were scaled to be bounded between 0 and 1, and inverted such that lower values indicate poorer flight performance. We define this measure as the flight performance index, which was analysed using a GLMM with a beta error distribution (*family = beta_family()* in R syntax).

Male mating success, defined as whether or not a mating occurred within 120 minutes, was analysed using a GLMM with a binomial error distribution.

For all feeding and flight response variables, the initial models included the factors sex, mating status, and dietary treatment (ethanol-supplemented vs. control), and all interaction terms. Block and Observer were included as random effects to account for variation due to experimental design and observer-related bias. For male mating response variables, initial models included male age as a continuous variable, dietary treatment (ethanol-supplemented vs. control), and their interaction term. Block and Observer were again included as random effects.

For all traits, model simplification was conducted by sequentially removing non-significant interaction terms, guided by likelihood ratio tests (via *anova;* Table S1-S5). Models with fewer parameters were retained when ΔAIC < 2 and further supported by lower BIC values, indicating a better balance between model fit and parsimony. After model selection, no interaction terms were retained in any of the final models, with the exception of male mating success. Diagnostics of all most parsimonious models were conducted using the *DHARMa* package (Hartig, 2025) to confirm normality, homoscedasticity, and appropriate dispersion of the model residuals. Significance of fixed effects was evaluated using Type III Wald chi-square tests via the Anova function from the *car* package (Fox & Weisberg, 2019). Post-hoc testing and estimation of marginal means were conducted using the *emmeans* package (Lenth & Piaskowski, 2025), and model and raw data visualisation was performed using the *ggplot2* package (Wickham, 2016).

## Results

Using a species whose adults naturally feed on decaying fruits, which can contain high levels of fermentation-derived ethanol (Dudley, 2004), we tested the hypothesis that ethanol consumption influences fitness-related behavioural traits. Specifically, through a series of behavioural assays, we examined the effects of dietary ethanol on feeding, flight endurance, and mating success, and assessed how these responses interact with individual status variables such as sex, mating status, and, when relevant, age.

### Dietary ethanol affects feeding and flight behaviours

We examined the effects of dietary ethanol, as well as sex and mating status on feeding-related variables (feeding latency and total feeding duration) and on a flight-related variable (flight performance).

All three predictor variables— sex, (χ² = 5.53, p = 0.018) mating status (χ² = 3.90, p = 0.048), and dietary ethanol treatment (χ² = 14.69, p < 0.001) — had a significant effect on feeding latency (Figure 1b; Table S1). On average, compared to females, males took approximately 50% longer to initiate feeding (females: 89.15 s [68.81–115.51], males: 133.81 s [102.78–174.22], mean [95%CI]). Mated individuals started feeding before virgin individuals (mated: 92.01 s [70.65–119.84], virgin: 129.65 s [99.99–168.11]), and individuals provided with an ethanol-containing diet initiated feeding before those given a control, non-ethanol diet; the latter taking approximately twice as long (experimental diet: 78.28 s [60.51–101.26], control diet: 152.41 s [116.72–199.00]).

**Figure 1.**
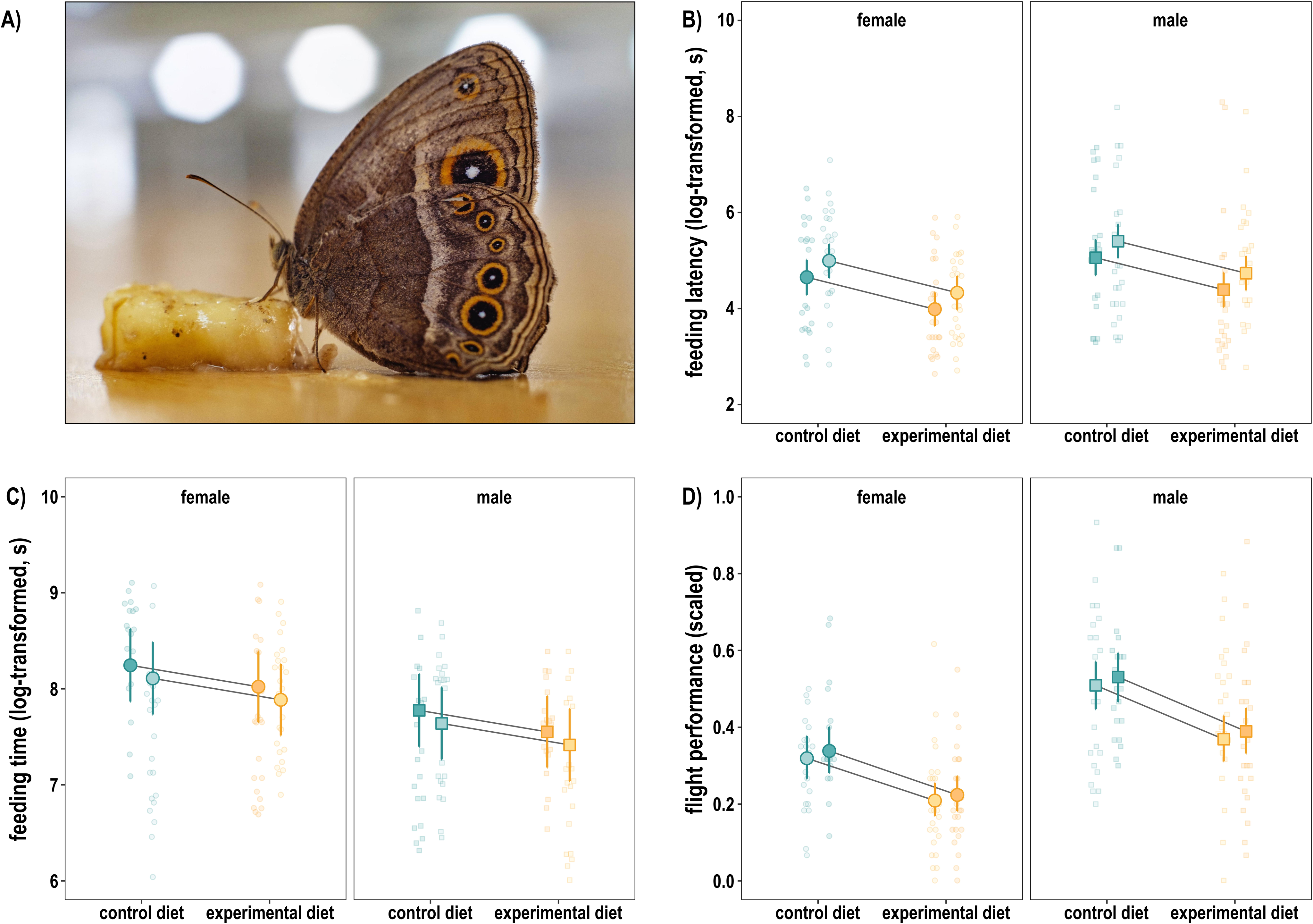
Effects of ethanol on feeding and flight behaviour. A) *Bicyclus anynana* female feeding on a slice of banana in the laboratory (image courtesy of Lin Zhaowei, reproduced with permission). B) Feeding latency of *B. anynana* was affected by dietary ethanol (*p* < 0.001), sex (*p* = 0.018) and mating status (*p* = 0.048), with females, mated individuals, and those on an ethanol-supplemented diet began feeding earlier than their respective controls. B) Total feeding time was affected by dietary ethanol (*p* < 0.030) and sex (*p* < 0.001); females fed longer than males, and individuals consuming an ethanol-supplemented diet fed shorter than those given a control diet. C) Flight performance was affected by dietary ethanol and sex (both: *p* < 0.001), with females and individuals on an ethanol-supplemented diet showing reduced flight performance. Details of statistical tests are provided in Tables S1-S3. B-D) Larger symbols represent model-estimated marginal means and smaller jittered symbols show raw values for all individuals; squares represent males and circles females; teal indicates the control diet and yellow the experimental diet; light-filled symbols denote unmated individuals and dark-filled symbols denote mated individuals. Error bars indicate 95% confidence intervals and include variability introduced by the random effects.

Total feeding time was significantly influenced by the sex of the individual (χ² = 21.68, *p* < 0.001) and the dietary ethanol treatment (χ² = 4.73, *p* < 0.030), but not by mating status (χ² = 1.67, *p* = 0.196) (Figure 1c; Table S2). On average, females fed for approximately 53.05 [37.61–74.83] minutes, which was about 60% longer than males (33.17 min [23.48–46.86]). Individuals consuming an ethanol- containing diet fed for approximately 9 minutes less, on average, than those on a non-ethanol diet (experimental diet: 37.51 min [26.64–52.82], control diet: 46.91 min [33.11–66.44]).

Both sex (χ² = 46.11, *p* < 0.001) and dietary ethanol treatment (χ² = 24.06, *p* < 0.001), but not mating status (χ² = 0.55, *p* = 0.456), had a significant effect on butterfly flight performance (Figure 1d; Table S3). On average, the flight performance index of males was higher than that of females (males: 0.45 [0.40–0.50], females: 0.27 [0.23–0.31]), indicating that males exhibited longer flight bouts than females. Individuals that consumed an ethanol-containing diet had a poorer flight performance compared to control individuals (experimental diet: 0.29 [0.25–0.33], control diet: 0.42 [0.37–0.47]).

### Dietary ethanol affects male mating behaviour

Behavioural assays were conducted to evaluate the effects of dietary ethanol on male mating success, and on mating latency among successful males.

For male mating success, we detected a significant interaction between male age and the diet consumed prior to the assay (Age × Diet: χ² = 7.28, p = 0.007). Post hoc testing revealed that dietary ethanol significantly reduced mating success of young males; males feeding on the control diet had over five times higher odds of mating compared to those feeding on the diet supplemented with ethanol (e.g., Age 4: odds ratio = 5.87, *z* = 3.24, *p* = 0.001). In contrast, no significant effect of diet was observed in older males (e.g., Age 10: odds ratio = 0.61, *z* = –1.15, *p* = 0.252), indicating that the detrimental impact of dietary ethanol on mating success diminishes with male age (Figure 2b; Table S4).

**Figure 2.**
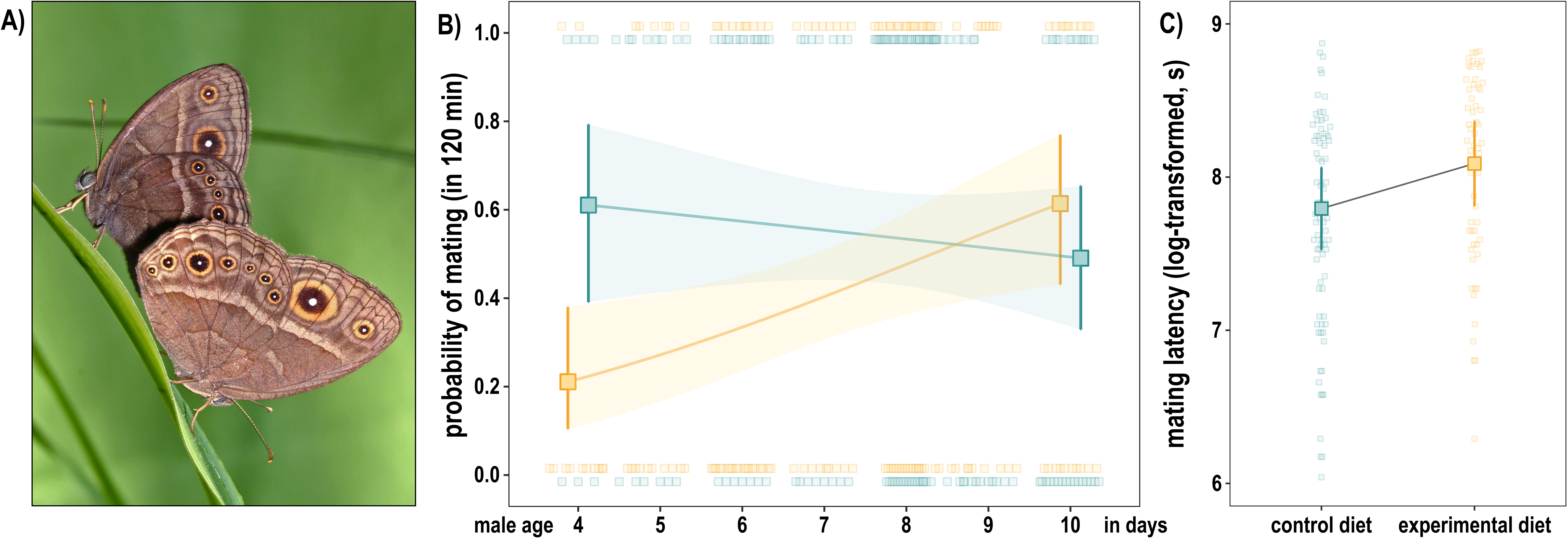
Effects of ethanol and age on mating success and mating latency. A) *Bicyclus anynana* male (top) mating with female (bottom) in the laboratory (image courtesy of William Piel, reproduced with permission). B) Male mating success was affected by an age-by-diet interaction (*p* = 0.007). Solid lines show model-predicted probabilities of male mating success as a function of age for both dietary treatments. Post hoc slope analysis revealed that age had no effect on the mating success of males consuming control diets (b = -0.08, [-0.30–0.14], *p* = 0.467), whereas it significantly increased the success of males receiving experimental diets (b = 0.30, [0.08–0.52], *p* = 0.008). Consequently, pairwise contrasts, represented by the larger symbols, revealed that dietary ethanol significantly reduced mating success in young males (4 days old; *p* = 0.001) but had no significant effect in older males (10-days old; *p* = 0.252). C) Mating latency of successful males in seconds was affected by dietary ethanol (*p* = 0.005) but not by male age (*p* = 0.762). On average, males consuming an ethanol-supplemented diet mated earlier than those given the control diet. B-C) Larger symbols represent model-estimated marginal means and smaller jittered symbols show raw values for all males, with teal representing the control diet and yellow the experimental diet. Shaded ribbons and error bars indicate 95% confidence intervals and include variability introduced by the random effects. Details of statistical tests are provided in Tables S4-S5.

Furthermore, mating latency of successful males was also significantly affected by dietary ethanol (χ² = 7.712, *p* = 0.005), but not by male age (χ² = 0.091, *p* = 0.762). On average, males feeding on the ethanol-containing diet took 54.26 [41.25–71.38] minutes to initiate mating, approximately 14 minutes longer compared to males feeding on a non-ethanol diet (40.45 min [30.99–52.79]; Figure 2c; Table S5).

## Discussion

### Effects of ethanol on feeding behaviour

In our study, *B. anynana* butterflies presented with ethanol-supplemented banana initiated feeding significantly faster than those offered the control diet (Fig 1B, Table S1). These findings align with previous feeding response assays, which demonstrated that *B. anynana* individuals consistently uncoiled their proboscis when exposed to solutions containing more than 1% ethanol (Dierks & Fischer, 2008; Kehl & Fischer, 2012). Ethanol-mediated foraging responses are widespread among nectar- and fruit-feeding animals (Bowland et al., 2024; Campbell et al., 2022; Sánchez et al., 2004) and their convergent evolution underscores both the reliability of ethanol as an olfactory cue and the significance of foraging-related behavioural traits for efficiently locating fermenting fruits in complex environments (Dudley, 2004).

We found that butterflies located the ethanol-supplemented diets more rapidly but spent less time feeding on this food (Fig 1C, Table S2). Reduced food intake at elevated dietary ethanol levels has been observed in other fruit-feeding butterflies (Reichert, 2011), hummingbirds (Choi et al., 2023), and fruit bats (Korine et al., 2011), highlighting that ethanol affects feeding behaviour across diverse taxa. It is, however, unclear whether the reduced feeding time in our experiments was observed because dietary ethanol impairs the butterflies’ ability to sustain feeding and/or because ethanol leads to faster satiation. The reduction in total feeding time could be a direct consequence of the physiological effects of ethanol and its primary metabolite, acetaldehyde, or of ethanol altering the palatability of the diet. On the other hand, enzymatic breakdown of ethanol via the alcohol detoxification pathway may allow these butterflies to use ethanol as an energy source (Bouchebti et al., 2024; Pohl et al., 2012), thereby diminishing the necessity for extended feeding, which can potentially reduce exposure to predators associated with terrestrial foraging (Young, 1979). Such utilisation of ethanol for energy and biosynthesis has been demonstrated in *Caenorhabditis* nematodes, where radiolabelled ethanol was fully oxidized to ¹⁴CO₂ (Cooper & Van Gundy, 1971), and L1 larvae incubated with low concentrations of ethanol exhibited an extended lifespan under starvation conditions (Castro et al., 2012). Consistent with these results, ethanol-supplemented diets were found to increase fecundity and survival in female *B. anynana* (Beaulieu et al., 2017), suggesting that ethanol can provide benefits for both reproduction and somatic maintenance.

Our feeding assays further revealed that females and mated individuals initiated feeding significantly faster than males and unmated individuals, respectively, and that females fed for longer than males. These results corroborate the established view that nutritional demands are key drivers of foraging and feeding behaviour (Boggs, 1992). In *B. anynana*, females are significantly larger than males (e.g. Rodrigues et al., 2021; van Bergen et al., 2017) and, thus, may have greater nutritional deficits following the experimental fasting period. Moreover, mated females require substantial nutrients for egg production, as stable isotope evidence demonstrated that approximately half of the carbon in *B. anynana* eggs is derived from the adult diet (Fischer et al., 2004). Nutritional demands may also differ between unmated and mated males, as the latter must replenish resources to produce a new spermatophore—a nutrient-rich nuptial gift that is energetically costly to produce (Boggs & Gilbert, 1979). These later-produced spermatophores have been shown to be approximately 25% smaller when *B. anynana* males are deprived of adult feeding (Ferkau & Fischer, 2006).

While our study found clear effects of dietary ethanol on feeding latency and duration, a previous study using the same species reported no significant effects on these behaviours (Beaulieu et al., 2017). This discrepancy may be attributed to differences in experimental design and procedures. In the earlier study, butterflies were fed ethanol-supplemented sucrose solutions, and behavioural assays were conducted with butterflies housed individually in one-litre transparent plastic containers, which restrict movement and natural foraging behaviour. In contrast, in our study, butterflies were assayed at low densities in hanging cages that allowed for free flight. In *B. anynana*, behavioural differences have been demonstrated to be more pronounced—and more readily detectable under experimental conditions—when butterflies are observed in free-flight environments compared to more restrictive laboratory settings (Joron & Brakefield, 2003; van Bergen & Beldade, 2019).

### Effects of ethanol on flight and male mating

In butterflies, effective flight underpins key behaviours, including foraging, predator avoidance, and mate acquisition, and, as such, it is critical for survival and reproduction (DeVries et al., 2010). Our results show that ethanol consumption impairs flight in *Bicyclus* butterflies (Fig 1D, Table S3). Moreover, males exhibited better flight performance in our experimental setup, likely reflecting sexual dimorphism in wing size (van Bergen et al., 2024), wing shape and associated aerodynamic parameters such as wing loading and aspect ratio (Saastamoinen et al., 2012), and flight muscle allocation (Saastamoinen et al., 2013). Although the dietary ethanol affected both males and females similarly in terms of flight, we focused on males, whose ritualized courtship involves flight and coordinated wing movements, to investigate the immediate fitness consequences of ethanol-induced flight impairment. Specifically, we compared the mating success of males that consumed an ethanol-supplemented diet with those that consumed a control diet. We found that diet-derived ethanol reduces the mating success of young males, and that this negative effect diminishes with male age (Fig 2B, Table S4). Moreover, ethanol consumption increased the time required for males to achieve a successful mating (Fig 2C, Table S5). To our knowledge, this is the first evidence that diet-derived ethanol can reduce male mating success in fruit-feeding Lepidoptera; however, Miller (1977) reported that adult fertility in *Choristoneura fumiferana* moths declined progressively with increasing ethanol concentrations, possibly as a result of reduced mating (Miller, 1997).

Male age is an important determinant of mating success in *Bicyclus* butterflies, with older males generally outcompeting younger ones (Fischer et al., 2008; Nieberding et al., 2012). The factors underlying this older male mating advantage are debated (Fischer et al., 2018; Nieberding & Holveck, 2018) but centre on two alternative hypotheses: (i) older males have lower residual reproductive value, which promotes more aggressive and risk-prone mating behaviour (Fischer et al., 2008), and (ii) females prefer the sex pheromone blend produced by older males (Nieberding et al., 2012). Ethanol consumption may influence male mating success by modulating mating motivation and/or pheromone composition in an age-dependent manner. For example, young males with high residual reproductive value may be more risk-averse under ethanol intoxication, whereas older males continue to seize mating opportunities as they arise. On the other hand, ethanol may provide precursors to produce the pheromones that play an important role in courtship. Consistent with this hypothesis, studies in fruit-feeding *Drosophila* have shown that attraction to dietary alcohol is particularly strong in unmated males (Shohat-Ophir et al., 2012), and that exposure to ethanol- and methanol-containing fruits results in a rapid increase in fatty acid–derived male sex pheromones (Keesey et al., 2025).

### Ecological and evolutionary significance of dietary ethanol

Shifts toward exploiting fruits as a primary nutritional resource required a suite of adaptations that allow animals to locate, consume, and process this complex food source (Herrera, 2002; Jordano, 2000). These adaptations extend beyond foraging behaviour and the physiological mechanisms that help mitigate the effects of ethanol in decaying fruits. For example, a key morphological innovation associated with fruit-feeding ecology is the serrated ovipositor of *Drosophila suzukii*, which enables the species to lay eggs on intact, ripe fruits (Akutsu & Matsuo, 2023) and allows the species to colonize the resource earlier than most other fruit-feeding competitors. This key morphological adaptation was accompanied by evolutionary change in other traits, including feeding behaviour (e.g. preference for hardness and shape characteristic of ripe fruit; Akutsu & Matsuo, 2022; Silva-Soares et al., 2017) as well as traits unrelated to feeding (e.g. genital morphology; Muto et al., 2018). Indeed, the evolution of frugivory may also affect traits not directly associated with resource acquisition. For instance, Young (1979) hypothesized that feeding on fallen, fermenting fruits increases exposure to predators — particularly when also intoxicated by ethanol — may have favoured the evolution of predator-avoidance signals (Young, 1979), such as conspicuous eyespot markings on the wings of various butterflies which are well-studied in *Bicyclus* (e.g., Beldade & Monteiro, 2021; Brattström et al., 2020). This perspective underscores how dietary shifts toward fruit feeding may shape the evolution of a wide range of traits, linking ecological specialization with evolutionary innovation, and highlighting the potential for fruit-derived compounds, such as ethanol, to influence physiology, behaviour, and fitness.

Effects of ethanol on animal health and fitness are typically dose-dependent: low doses can be beneficial, whereas higher doses are often harmful or toxic—a biological phenomenon known as hormesis. For example, low doses of dietary ethanol can increase the survival of Drosophila larvae (Schumann et al., 2021), including by providing protection against endoparasitoids (Milan et al., 2012), while high doses remain toxic and increase larval mortality (Schumann et al., 2021). The concentration used in our study (9.6% v/v) falls within the range employed by comparable experimental work (e.g. Bouchebti et al., 2024; Dierks & Fischer, 2008; Schumann et al., 2021), although it might exceed the typical levels that these butterflies encounter in natural settings. However, while most fruits accessible to frugivores in tropical habitats indeed contain relatively low ethanol concentrations, typically around 1% v/v (Casorso et al., 2023; Maro et al., 2025), overripe fruits can reach much higher levels; for example, ethanol concentrations as high as 10.3% v/v have been detected in *Astrocaryum standleyanum* palm fruits in Panama (Dudley, 2004). The high concentration in our study may have accentuated harmful or toxic effects, while overlooking potential beneficial effects present at lower concentrations. However, it did uncover a range of effects which invite exciting avenues for further investigation. Beyond investigating dose-dependent effects, future studies could explore ethanol effects on male-male competition and on female choice, as well as on the molecular mechanisms by which dietary ethanol and its metabolism might impact feeding, flight and male mating behaviours. These studies can shed light on the evolutionary forces driving ethanol tolerance in fruit-feeding butterflies and other frugivorous animals.

## Conclusion

We investigated the effects of dietary ethanol on feeding behaviour, flight performance and mating success in fruit-feeding *B. anynana* butterflies. Our results revealed that butterflies found ethanol-supplemented diets faster, but spent less time feeding on these diets. Moreover, ethanol consumption impaired overall flight performance and mating success of young males. Collectively, our findings suggest that tropical *Bicyclus* butterflies can use ethanol as an olfactory cue to locate suitable food sources in their natural environment (Molleman et al., 2005). Nonetheless, our study also clearly demonstrates that ethanol consumption alters feeding behaviour, flight performance and mating success of these fruit-feeding insects, despite *B. anynana* likely possessing effective detoxification mechanisms (Conceição et al., 2011; Nowell et al., 2017). Our results point to a potential cost–benefit trade-off associated with diet-derived ethanol in natural contexts, and thus contribute to ongoing efforts to understand the ecological and evolutionary significance of ethanol consumption across animal lineages.

## Acknowledgements

We are grateful to Ana Rita Garrizo, Guilherme Atencio and Carlyne Golding for technical assistance. Financial support provided by the Portuguese science funding agency, *Fundação para a Ciência e Tecnologia* (FCT; grant IDs: PTDC/BIA-EVL/0321/2021 and 2023.18093.ICDT) to PB. SH was funded by the European Commission through Marie Skłodowska-Curie Actions fellowship (grant ID: 101104682). EvB received salary support through European Research Council (grant ID: 101042392, awarded to Inês Fragata).

