## Supplementary Materials for "Dietary ethanol affects feeding behaviour, flight performance, and mating success in a fruit-feeding butterfly"


| | |
|---|---|
| Supplementary Fig. 1 | Schematic representation of the experimental design. |
| Supplementary Table 1 | Generalized linear mixed models for feeding latency. |
| Supplementary Table 2 | Generalized linear mixed models for total feeding time. |
| Supplementary Table 3 | Generalized linear mixed models for flight performance. |
| Supplementary Table 4 | Generalized linear mixed models for male mating success. |
| Supplementary Table 5 | Generalized linear mixed models for male mating latency. |

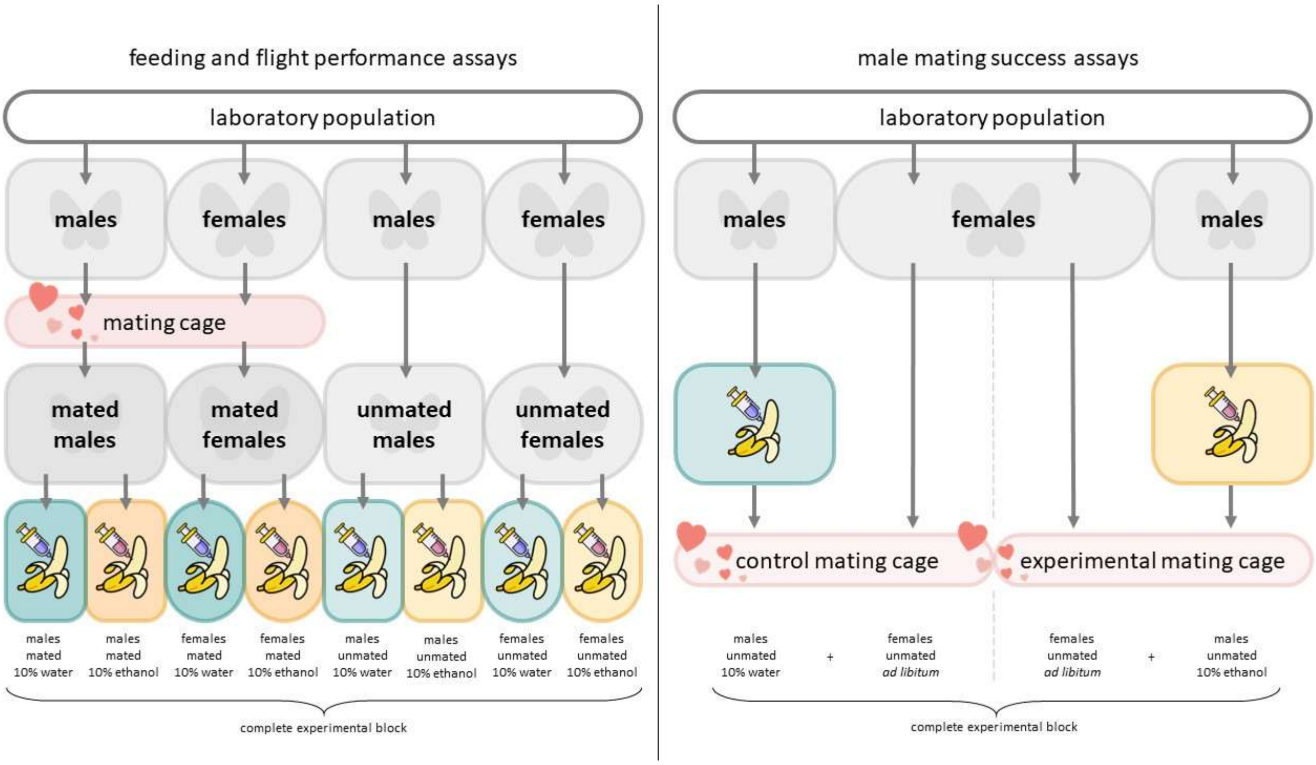


**Supplementary Figure 1.** Schematic representation of the experimental design. Panels on the left-hand side depict the design of the feeding and flight assays while panels on the right-hand side provide an overview of the male mating assays. For both assays, butterflies were reared at 27 °C. Pupae were collected from the laboratory population, transferred to individual containers, and individually marked as adults upon emergence. For the feeding and flight assays (left), individuals were first assigned to one of two mating treatments: half of the individuals of each sex were allowed to mate, while the other half remained unmated. After mating, individuals were transferred to separate cages corresponding to mated males, mated females, unmated males, and unmated females. After a brief fasting period, butterflies were further divided into two dietary treatments: a control diet (teal) or an experimental diet (yellow). Feeding and flight behaviour assays were conducted on a total of 200 individuals across five replicate blocks. For the male mating assays (right), virgin males were allocated to one of two cages, while all females were maintained together. After a brief fasting period, males were assigned to one of two dietary treatments: a control diet (teal) or an experimental diet (yellow). After males completed feeding, virgin females were added to each mating cage at a 1:1 sex ratio. Male mating success and mating latency of successful males were assessed using 295 males, aged three to ten days, across 16 replicate blocks.

**Table S1a.** Model selection for log-transformed feeding latency (in seconds) using GLMMs (glmmTMB, Gaussian). All models included random intercepts for Block and Observer. Starting from the full model with all interactions (Sex × State × Food; model 1), interactions were sequentially removed to identify the most parsimonious model (model 5), based on AICc and BIC.

| **feeding latency** | *Food* | *Sex* | *State* | *Food:Sex* | *Food:State* | *Sex:State* | *Food:Sex:State* | **df** | **logLik** | **AICc** | **ΔAICc** | **wAICc** | **BIC** | **ΔBIC** | **wBIC** |
|---|---|---|---|---|---|---|---|---|---|---|---|---|---|---|---|
| model 5 | + | + | + | | | | | 7 | -295.009 | 604.648 | 0.000 | 0.565 | 626.599 | 0.000 | 0.904 |
| model 4 | + | + | + | | + | | | 8 | -294.731 | 606.276 | 1.628 | 0.250 | 631.268 | 4.670 | 0.088 |
| model 3 | + | + | + | + | + | | | 9 | -294.530 | 608.083 | 3.435 | 0.101 | 636.092 | 9.493 | 0.008 |
| model 2 | + | + | + | + | + | + | | 10 | -294.530 | 610.316 | 5.669 | 0.033 | 641.317 | 14.718 | 0.001 |
| model 1 | + | + | + | + | + | + | + | 11 | -292.977 | 609.471 | 4.824 | 0.051 | 643.437 | 16.839 | 0.000 |

**Table S1b.** Results of the Type III Wald chi-square tests for the final GLMM (model 5) testing the effects of Sex, State (mating state), and Food (dietary ethanol treatment) on log-transformed feeding latency (in seconds).

| **Term** | **ChiSq** | **df** | **Pr(>ChiSq)** |
|---|---|---|---|
| (Intercept) | 633.3313 | 1 | < 0.001 |
| Sex | 5.5279 | 1 | 0.0187 |
| State | 3.9044 | 1 | 0.0482 |
| Food | 14.6919 | 1 | < 0.001 |

**Table S2a.** Model selection for total feeding time (in seconds) using GLMMs (glmmTMB, Gamma family with log link). All models included random intercepts for Block and Observer. Starting from the full model with all interactions (Sex × State × Food; model 1), interactions were sequentially removed to identify the most parsimonious model (model 5), based on AICc and BIC.

| feeding duration | *Food* | *Sex* | *State* | *Food:Sex* | *Food:State* | *Sex:State* | *Food:Sex:State* | df | logLik | AICc | ΔAICc | wAICc | BIC | ΔBIC | wBIC |
|---|---|---|---|---|---|---|---|---|---|---|---|---|---|---|---|
| model 5 | + | + | + | | | | | 7 | -1572.868 | 3160.383 | 0.000 | 0.513 | 3182.125 | 0.000 | 0.887 |
| model 4 | + | + | + | | | + | | 8 | -1572.430 | 3161.697 | 1.314 | 0.266 | 3186.448 | 4.322 | 0.102 |
| model 3 | + | + | + | | + | + | | 9 | -1572.145 | 3163.343 | 2.960 | 0.117 | 3191.077 | 8.952 | 0.010 |
| model 2 | + | + | + | + | + | + | | 10 | -1572.061 | 3165.416 | 5.033 | 0.041 | 3196.107 | 13.981 | 0.001 |
| model 1 | + | + | + | + | + | + | + | 11 | -1570.516 | 3164.593 | 4.210 | 0.063 | 3198.215 | 16.090 | 0.000 |

**Table S2b.** Results of the Type III Wald chi-square tests for the final GLMM (model 5) testing the effects of Sex, State (mating state), and Food (dietary ethanol treatment) on total feeding time (in seconds).

| Term | ChiSq | df | Pr(>ChiSq) |
|---|---|---|---|
| (Intercept) | 1838.0310 | 1 | < 0.001 |
| Sex | 21.6834 | 1 | < 0.001 |
| State | 1.6705 | 1 | 0.1962 |
| Food | 4.7276 | 1 | 0.0297 |

**Table S3a.** Model selection for flight performance using GLMMs (glmmTMB, Beta family). All models included random intercepts for Block and Observer. Starting from the full model with all interactions (Sex × State × Food; model 1), interactions were sequentially removed to identify the most parsimonious model (model 5), based on AICc and BIC.

| **flight performance** | *Food* | *Sex* | *State* | *Food:Sex* | *Food:State* | *Sex:State* | *Food:Sex:State* | **df** | **logLik** | **AICc** | **ΔAICc** | **wAICc** | **BIC** | **ΔBIC** | **wBIC** |
|---|---|---|---|---|---|---|---|---|---|---|---|---|---|---|---|
| model5 | + | + | + | | | | | 7 | 72.354 | -130.052 | 0.000 | 0.539 | -108.395 | 0.000 | 0.887 |
| model4 | + | + | + | + | | | | 8 | 72.776 | -128.705 | 1.347 | 0.275 | -104.053 | 4.342 | 0.101 |
| model3 | + | + | + | + | | + | | 9 | 73.101 | -127.138 | 2.914 | 0.125 | -99.516 | 8.879 | 0.010 |
| model2 | + | + | + | + | + | + | | 10 | 73.215 | -125.121 | 4.931 | 0.046 | -94.557 | 13.839 | 0.001 |
| model1 | + | + | + | + | + | + | + | 11 | 73.238 | -122.895 | 7.157 | 0.015 | -89.415 | 18.980 | 0.000 |

**Table S3b.** Results of the Type III Wald chi-square tests for the final GLMM (model5) testing the effects of Sex, State (mating state), and Food (dietary ethanol treatment) on log-transformed feeding latency (in seconds).

| **Term** | **ChiSq** | **df** | **Pr(>ChiSq)** |
|---|---|---|---|
| (Intercept) | 23.6228 | 1 | < 0.001 |
| Sex | 46.1054 | 1 | < 0.001 |
| State | 0.5546 | 1 | 0.4564 |
| Food | 24.0580 | 1 | < 0.001 |

**Table S4.** Results of the Type III Wald chi-square tests for a GLMM (glmmTMB, binomial family) testing the effects of Food (dietary ethanol treatment), and Age (days since eclosion), and their interaction (Age × Food) on male mating success. Model included random intercepts for Block and Observer.

| **Term** | **ChiSq** | **df** | **Pr(>ChiSq)** |
|---|---|---|---|
| (Intercept) | 0.7843 | 1 | 0.3758 |
| Age | 0.5296 | 1 | 0.4668 |
| Food | 9.3304 | 1 | 0.0023 |
| Age:Food | 7.2787 | 1 | 0.0070 |

**Table S5a.** Model selection for mating latency (in minutes) using GLMMs (glmmTMB, Gamma family with log link). All models included random intercepts for Block and Observer. Starting from the full model with all interactions (Age × Food; model 1), interactions were sequentially removed to identify the most parsimonious model (model 2), based on AICc and BIC.

| **mating latency** | *Age* | *Food* | *Age:Food* | **df** | **logLik** | **AICc** | **ΔAICc** | **wAICc** | **BIC** | **ΔBIC** | **wBIC** |
|---|---|---|---|---|---|---|---|---|---|---|---|
| model 2 | + | + | | 6 | -1216.324 | 2445.288 | 0.000 | 0.743 | 2462.211 | 0.000 | 0.918 |
| model 1 | + | + | + | 7 | -1216.273 | 2447.407 | 2.119 | 0.257 | 2467.036 | 4.826 | 0.082 |

**Table S5b.** Results of the Type III Wald chi-square tests for the final GLMM (model 2) testing the effects of Age and Food (dietary ethanol treatment) on mating latency (in minutes).

| **Term** | **ChiSq** | **df** | **Pr(>ChiSq)** |
|---|---|---|---|
| (Intercept) | 816.2420 | 1 | < 0.001 |
| Age | 0.0914 | 1 | 0.7624 |
| Food | 7.7124 | 1 | 0.0055 |
